# Sequences at gene segment termini inclusive of untranslated regions and partial open reading frames play a critical role in mammalian orthoreovirus S gene packaging

**DOI:** 10.1101/2023.05.25.542362

**Authors:** Debarpan Dhar, Samir Mehanovic, Walter Moss, Cathy L. Miller

**Affiliations:** Interdepartmental Microbiology Graduate Program, Iowa State University, Ames, Iowa, USA; Department of Veterinary Microbiology and Preventive Medicine, College of Veterinary Medicine, Iowa State University, Ames, Iowa, USA; Roy J. Carver Department of Biochemistry, Biophysics and Molecular Biology, Iowa State University, Ames, Iowa, USA

## Abstract

Mammalian orthoreovirus (MRV) is a prototypic member of the *Spinareoviridae* family and has ten double-stranded RNA segments. One copy of each segment must be faithfully packaged into the mature virion, and prior literature suggests that nucleotides (nts) at the terminal ends of each gene likely facilitate their packaging. However, little is known about the precise packaging sequences required or how the packaging process is coordinated. Using a novel approach, we have determined that 200 nts at each terminus, inclusive of untranslated regions (UTR) and parts of the open reading frame (ORF), are sufficient for packaging each S gene segment (S1-S4) individually and together into replicating virus. Further, we mapped the minimal sequences required for packaging the S1 gene segment to 25 5′ nts and 50 3′ nts. The S1 UTRs alone are not sufficient, but are necessary for packaging, as mutations of the 5′ or 3′ UTRs led to a complete loss of virus recovery. Using a second novel assay, we determined that 50 5′nts and 50 3′ nts of S1 are sufficient to package a non-viral gene segment into MRV. The 5′ and 3′ termini of the S1 gene are predicted to form a panhandle structure and specific mutations within the predicted stem of the panhandle region led to a significant decrease in viral recovery. Additionally, mutation of six nts that are conserved in the three major serotypes of MRV and are predicted to form an unpaired loop in the S1 3′UTR, led to a complete loss of viral recovery. Overall, our data provide strong experimental proof that MRV packaging signals lie at the terminal ends of the S gene segments and offer support that the sequence requirements for efficient packaging of the S1 segment include a predicted panhandle structure and specific sequences within an unpaired loop in the 3′ UTR.

## INTRODUCTION

The process of gene segment packaging, whereby one copy of each of multiple gene segments are faithfully incorporated into a full genome, a virion is assembled, and the genome is replicated remains a persistent mystery in our understanding of the basic biology of segmented dsRNA viruses in the Order *Reovirales*. Mammalian orthoreovirus (MRV) has long been studied as a prototypical member of this Order which also includes important animal and human pathogens such as bluetongue virus (BTV) and rotavirus (RV). MRV is considered clinically benign, however it is a potent oncolytic virus that has been widely studied as a cancer therapeutic in Phase I-III clinical trials and has recently been awarded FDA fast track designation for breast and pancreatic cancer therapy (1–3). Therefore, it is critical to resolve remaining questions in the MRV life cycle to both answer longstanding important biological questions and provide actionable insight into its development as a cancer therapeutic.

MRV has ten double stranded (ds)RNA genome segments [three large (L) encoding, three medium (M), and four small (S)] that encode for twelve proteins. Eight of the twelve proteins form the bilayered virus capsid, within which the ten dsRNA genome segments are packaged (4). During genome packaging, viruses with multi-segmented genomes must coordinate the faithful incorporation of one copy of each gene segment into the virion. This may be achieved through two fundamental strategies, one where the gene segments are “fed” into a preassembled empty viral capsid and another where the gene segments act as the scaffold around which the viral proteins assemble (5–7). For φ6 bacteriophage of the *Cystoviridae* family that have a genome of three dsRNA segments, each genome segment is fed in a specific order (Small (s+)>Medium (m+)>Large (l+)) into an assembled procapsid by a helicase-dependent mechanism. The packaging process is dually dependent on a conserved 18 nt sequence stretch present in all three segments and unique packaging (pac) sequences located at the 5′ end of each of the segments (8–10). During φ6 packaging, the s+ gene pac sequence binds to a specific site on the procapsid vertices where it is then translocated into the procapsid. Packaging of the s+ segment results in conformational changes and expansion of the procapsid that reveal recognition sites necessary for packaging of the m+ segment, and following additional procapsid expansion, the l+ segment.

For such a mechanism to work in dsRNA viruses with larger genomes (9-12 segments), it would necessitate up to 12 conformational changes of the preformed core to package the RNA (6). As empty core particles are structurally indistinct from those containing a full genome, it is unlikely that members of the *Spinareoviridae* or *Sedoreoviridae* families use such a mechanism for packaging. Instead, evidence suggests that for segmented dsRNA viruses with larger genomes, packaging is likely guided through RNA:RNA interactions between the plus-strand RNAs of each segment. For example, *in vitro* studies revealed that RNA interactions between the ten plus-strand RNA segments of BTV are essential for assembling the viral core (11), and genome packaging is a sequential process, likely triggered by the UTR of the smallest gene segment (S10) that subsequently recruits the medium and larger segments via RNA:RNA interactions to form an assortment complex (12). For RV, capsid proteins assemble around a plus-strand RNA/capping enzyme/RNA-dependent RNA Polymerase (RdRP) complex. This assembly is likely enhanced and stabilized by the interactions of the capsid protein N terminal extensions with the +ssRNAs (13). RV plus-strand RNA segments also have been shown to interact with each other by RNA:RNA interactions. Packaging likely follows a sequential order with the smaller S segments initiating the process. In addition, there is a reduction in viral complex formation and replication when the UTRs are targeted using sequence-specific antisense oligoribonucleotides (ORNs)(14).

Another piece of evidence that suggests that MRV probably uses a selective packaging mechanism is that the ratio of total particles to infectious particles for MRV is low (< 2) (15, 16), indicating that the MRV packaging mechanism efficiently produces progeny virions with an entire complement of genes. In addition, structural studies on MRV and other dsRNA viruses have shown that the number of transcription complexes correlates with the number of gene segments, despite structural space for additional complexes (17–20). Altogether, these data lend support that MRV uses a selective genome packaging mechanism for producing progeny virions.

The selective packaging mechanism of MRV can be imagined as a process made up of three fundamental processes. First is *Assortment*, where the individual plus-strand RNAs for all ten genome segments (S1-S4, M1-M3, and L1-L3) interact to form a sorted set of the full genome. *Assembly* of the inner capsid proteins to form the viral core either quickly follows or coincides with assortment. Recent evidence suggests that a single-layered particle (SLP) may form as an assembly intermediate, and that the assortment complex is brought to the SLP by the RdRp (λ3), which is bound to each gene segment terminus (21, 22). Second strand/negative-strand synthesis (*Replication*) of the plus-strand RNAs to dsRNAs by λ3 and a polymerase co- factor μ2 is thought to occur during or shortly after viral core assembly (4).

MRV genomes have short UTRs that range from 12-32 nts (5′ end), and 35-83 nts (3′ end), with a string of 4 (5′ GCUA) and 5 (3′ UCAUC) nts at the termini of each segment that are conserved in all serotypes (23–26). Similar terminal short conserved sequences are critical for RV polymerase binding, suggesting their primary function is in replication, and not assortment/assembly (27). Moreover, defective interfering particle gene segments with large internal gene deletions always retain the UTRs and variable lengths of the ORF at each end (26, 28), suggesting that packaging signals extend beyond the UTRs into the ORF. Based on published data from MRV and other viruses with a dsRNA genome (26, 28–33), we have hypothesized that signal sequences present at the terminal ends of each MRV gene segment inclusive of UTRs and partial ORFs guide the overall process of packaging (assortment, assembly, and replication).

As a result of the extension of packaging signals into the ORF, and the dependence of successful MRV progeny production on the expression of all viral proteins, identifying packaging signals experimentally has proven challenging. Initial experiments to identify MRV packaging sequences relied on a complicated reverse genetics (RG) system in which a plasmid encoding the chloramphenicol acetyltransferase (CAT) gene was flanked on each end with MRV gene segment terminal sequences, followed by recombinant virus rescue using a helper virus and cell lines expressing the protein encoded by the deleted ORF (29–32). More recently, a novel approach where MRV gene segment protein-coding regions were unlinked from putative packaging signals by duplicating terminal regions inclusive of the ORF and UTR, then introducing mutations in the third/wobble codon position in the ORF region was used successfully to identify sequences involved in MRV packaging. (33). The prior research was rigorous and generated significant novel information highlighting the importance of the terminal regions of the MRV genes towards packaging. However, these approaches were experimentally complex, relying on helper cell lines or the addition of duplicated sequences. Furthermore, in some cases these approaches could not rule out the possibility of wildtype (WT) sequences within the internal regions of the ORF contributing to the packaging process. Moreover, these approaches often did not identify minimal RNA sequences or structures required for the packaging process.

In this work we have created a novel approach, termed Wobble/ Block Replacement (W/BR), that uses codon redundancy to introduce the maximum possible nt changes across the varying regions of the ORF of each MRV gene segment without altering amino acid sequences. Using this approach, we have identified the importance of terminal nt sequences in the packaging of S gene segments (S1-S4) of the Type 1 Lang (T1L) serotype. Furthermore, for the first time, we have identified minimal sequences in the S1 gene that are sufficient for packaging. We have also provided experimental support that sequences within a predicted panhandle structure formed by the interaction of the terminal ends of the MRV gene play an important role in S1 packaging. Finally, we have used a novel Segment Incorporation Assay to investigate the sufficiency of packaging signal sequences in incorporating a non-viral reporter gene into infectious MRV. Our results are interesting from a fundamental biology standpoint in shedding light on the black box of dsRNA virus assortment. Additionally, because MRV is being investigated as an oncolytic virus, our findings may be of use clinically by identifying packaging sequences that are indispensable for the creation of MRV oncotherapeutic vectors.

## RESULTS

### Wobble/Block Replacement (W/BR) is a novel approach to identify MRV packaging sequences

A central problem in determining packaging sequences (PAC) within the MRV genome is that predicted sequences necessary for assortment/assembly/replication extend beyond the UTRs and overlap with the protein-coding region (Fig. 1A, Top). This makes the identification of packaging sequences challenging, as it is difficult to introduce mutations or deletions into gene segments without impacting MRV proteins. Our novel W/BR approach overcomes these limitations by taking advantage of codon redundancy to make maximum nt changes throughout the ORF (Fig. 1A, Bottom) while ensuring that the amino acid sequence remains unchanged. Using this approach, we designed S1-S4 mutant constructs (18/S1/40, 23/S2/59, 32/S3/73, 37/S4/69) where each gene contained WT sequence in both 5′ and 3′ UTRs as well as start and stop codons, but every possible nt within the ORF was “wobbled”. These “wobble” changes were not limited to nts in the third/wobble position but include double nt changes in leucine and arginine and triple nt changes in serine codons. These wobble mutant constructs have been named uniformly throughout the publication in the format X/Gene/Y, where X and Y denote the number of wild-type sequences at the 5′ and 3′ end, respectively, of a wobbled gene segment. As a result of the wobble changes within the ORF, the four S gene segments are only 57 to 61% identical to their corresponding wild-type gene sequences (Fig. 1A). Plasmids are also made in which terminal regions of varying lengths within the ORF contain wild-type sequences to identify packaging requirements at the sequence level. To determine packaging requirements, each of these constructs are subjected to the MRV reverse genetics (RG) assay and the impact of the mutations on recombinant virus recovery is determined by viral titer on a standard MRV plaque assay. Mutant plasmids expressing gene segments that contain nt sequences sufficient for packaging generate replicating viruses that form plaques on L929 cells, which are picked and passaged for amplification, viral RNA extraction, reverse transcription polymerase chain reaction (RT-PCR), and sequencing (Fig. 1B).

**Fig. 1.**
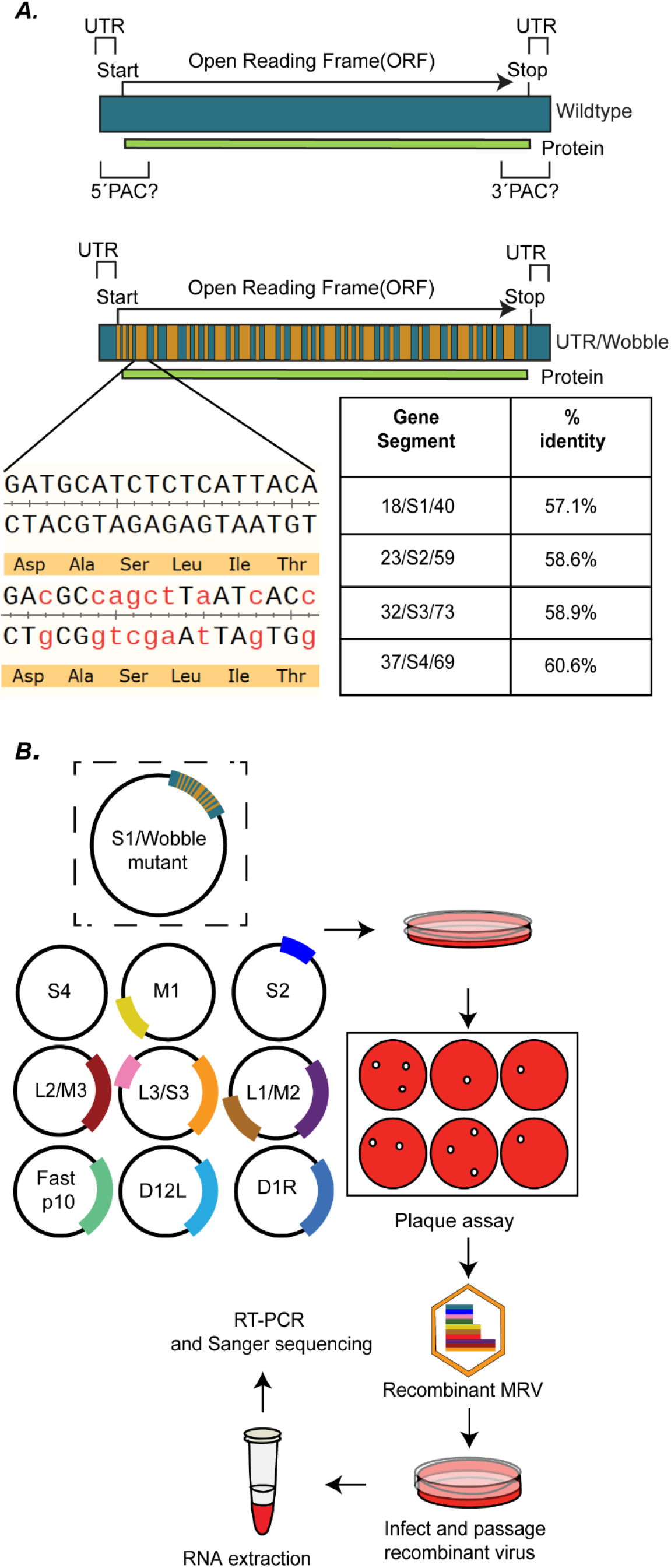
Wobble/Block Replacement is a novel approach to identify MRV packaging sequences. (A) (Top) Illustration of an MRV WT S gene segment showing short UTRs, and predicted packaging signals that extend beyond the UTR into the open reading frame. (Bottom) Illustration of wobble mutation strategy with WT nt regions represented in green, and wobble mutations represented in orange. An example of wobble mutations and the percent change to the nt sequences compared to WT are shown. (B) Schematic of the W/BR assay showing replacement of WT S1 with an S1 wobble mutant. The S1 wobble mutant is transfected with plasmids encoding for the remaining nine WT MRV gene segments and accessory plasmids into BHK-T7 cells in the MRV RG assay. Viral titer is determined using plaque assay, and isolated plaques are picked and passaged to amplify the viral titer before RNA extraction, RT-PCR amplification, and sequencing are completed.

### 200 WT nts but not UTRs at each terminus are sufficient for efficient recovery of replicating virus

To provide proof of concept of the W/BR assay, sets of two wobble mutant constructs were designed for each S gene segments, S1 (encoding σ1 and σ1s), S2 (encoding σ2), S3 (encoding σNS) and S4 (encoding σ3). In each set, one construct maintained WT sequences only at the terminal ends comprising the UTR, Start codon and 2 non-wobbled nts (5′ end) and 3′ UTR and Stop codons (3′ end) while the ORF was fully wobbled (18/S1/40, 23/S2/59, 32/S3/73, 37/S4/69). In the second construct in each set, 200 nt sequences at each termini, inclusive of the UTRs, Stop and Start codons and variable lengths of the ORF maintained WT nt sequences, while the rest of the ORF remained fully wobbled (200/S1/200, 200/S2/200, 200/S3/200, 200/S4/200) (Fig. 2A). Each set of mutant clones for S1, S2, S3 and S4 genes, as well as their WT gene counterparts were subjected to RG assays and viral titer of each RG lysate was determined by plaque assay on L929 cells. No plaques were formed following RG with 23/S2/59, 32/S3/73, or 37/S4/69 constructs, strongly suggesting the UTRs are insufficient to support the packaging of S2, S3 and S4 gene segments (Fig. 2B). For the S1 gene, the 18/S1/40 virus was recovered at levels 10,000-100,000 fold lower than the WT S1, and produced significantly smaller plaques than WT and 200/S1/200 (Fig. 2C), suggesting that while packaging was not completely lost with this plasmid, it was severely and significantly decreased. On the other hand, 200/S1/200, 200/S2/200, 200/S3/200, and 200/S4/200 plasmid constructs in RG supported recombinant virus production at levels not significantly different from the corresponding WT plasmids (Fig. 2B). These results indicate that when the ORF nt sequence is mutated substantially by wobble introduction, the UTRs are unable to support efficient packaging of the individual S genes. However, even though wobble mutations are maintained across the majority of the gene segment, reversion of 200 WT nts at each terminal end inclusive of the UTRs and parts of the ORF is sufficient for packaging the S gene segments individually into replicating viruses to WT levels.

**Fig. 2.**
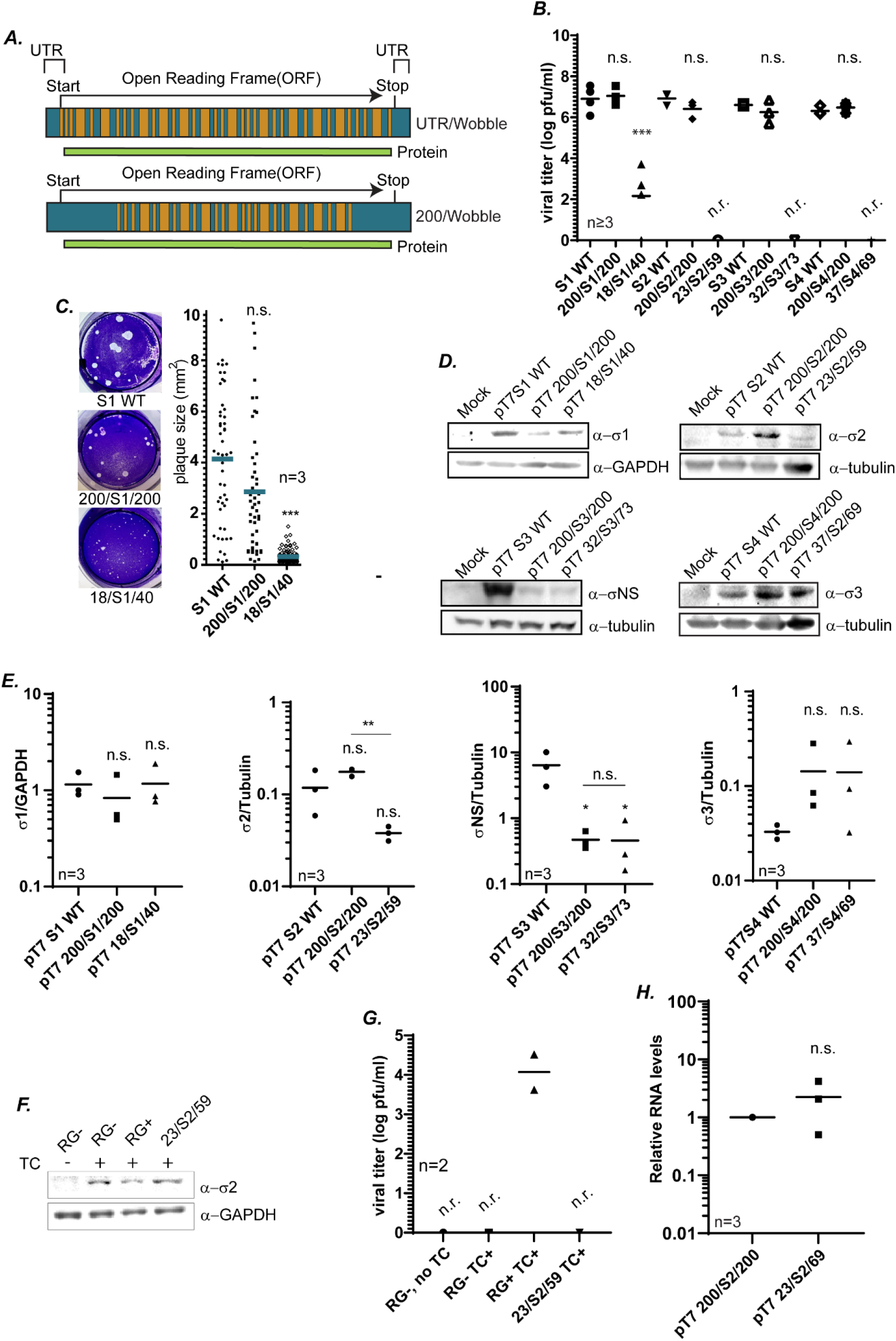
200 WT nt but not WT UTRs at each terminus are sufficient for efficient recovery of replicating virus. (A) Illustration of UTR/Wobble/UTR and 200/Wobble/200 constructs created for use in the W/BR assay. (B) Viral titers (log pfu/ml) of the S WT, 200/S gene/200, and UTR/S gene/UTR gene segments (S1-S4) measured after RG and plaque assay (C) Viral plaque size images (left) and quantifaction (right) of 200/S1/200, 18/S1/40, and WT S1 recombinant viruses recovered from RG. (D) Representative immunoblots using specific σ antibodies showing protein expression of σ proteins from indicated S gene plasmids 24 h post-transfection. (E) Quantification of biological replicates of immunoblots in (D). σ protein intensities are normalized to the corresponding loading controls (GAPDH, TUBULIN). (F) Immunoblots showing σ2 expression from RG lysates in the σ2 transcomplementation assay. (G) Viral titer (log pfu/ml) from RG lysates measured by plaque assay in the σ2 transcomplementation assay (H) Relative S2 RNA levels measured using RT-qPCR and normalized to housekeeping gene *β-actin* (*ACTB*). TC: Transcomplementation, RG-: Reverse genetics negative control (no pT7-S2). RG+: Reverse genetics positive control (pT7-S2 WT). One-way ANOVA and Tukey comparisons were used for statistical analyses of data, except for Fig. 2G where Student t-test was performed. n= independent biological replicates, n.r.=virus not recovered, *p<0.05, ** p<0.01, *** p<0.001, n.s.=non-significant, p>0.05.

Though the WT amino acid sequence of each gene was maintained upon introduction of wobble mutations, there is a possibility that defects in the overall protein expression of the σ proteins were altered in the RG assay, which could impact virus rescue. To examine this possibility, S gene segment wobble mutant plasmids were transfected into BHK-T7 cells and at 24 hours post transfection, cells were processed for immunoblotting to measure protein expression. Despite vastly different efficiencies in virus recovery, there were no significant differences in σ1 protein expression in pT7 S1 WT, pT7 200/S1/200 and pT7 18/S1/40 (Fig. 2D and 2E). When measuring σ2 expression, we noted a significant decrease in expression of the pT7 23/S2/59 construct relative to pT7 200/S2/200, although there was no significant difference from WT pT7 S2 (Fig. 2D and 2E). To determine whether a decrease in σ2 protein expression contributed to the loss of virus recovery of 23/S2/59 virus (Fig. 2B), we performed a transcomplementation assay. In this assay, RG was performed with the addition of a plasmid expressing the WT σ2 protein from a construct that lacks the T7 promoter and Hepatitis δ ribozyme, therefore, the RNA produced does not resemble the authentic S2 gene segment and cannot be incorporated in place of S2 into the MRV genome for virus rescue. The addition of this plasmid in the RG assay resulted in similar expression of σ2 across all samples (Fig. 2F) but did not result in 23/S2/59 virus recovery (Fig. 2G). This suggests that increasing σ2 protein expression in RG assays using pT7 23/S2/59 does not result in recovery of virus in the RG assay, and that the defect in recovery lies outside of protein expression. A second possibility that we considered for lack of RG recovery of 23/S2/59 was that of RNA expression. To determine whether the defect in viral recovery of the 23/S2/59 virus was due to a lack of S2 RNA available for packaging, we performed an RT-qPCR assay to measure the relative RNA levels of S2 (Fig. 2H). The accumulated RNA from the pT7 23/S2/59 and pT7 200/S2/200 construct after 24 hours of transfection was normalized to the housekeeping gene *β-actin (ACTB)*. The normalized 23/S2/59 RNA was compared to the reference construct of normalized 200/S2/200 RNA. Our results showed no significant changes in the RNA levels of 23/S2/59 compared to the 200/S2/200.

While 200/S3/200 virus is recovered to WT levels, the 32/S3/73 virus was not recovered (Fig. 2B), however pT7 200/S3/200 and pT7 32/S3/73 express σNS protein similarly with respect to each other (Fig. 2D and 2E), suggesting that protein expression levels are not playing a role in loss of 32/S3/73 recovery. However, it is interesting to note that while 200/S3/200 and S3 WT viruses were recovered from RG at similar levels (Fig. 2B), the pT7 S3 WT protein was expressed at significantly higher levels than either pT7 200/S3/200 or pT7 32/S3/73 (Fig. 2D and 2E). This has little bearing on our results but suggests that WT S3 expresses much higher levels of protein than are needed for efficient virus recovery. Similarly, we did not see significant differences in protein expression in immunoblots of σ3 between 200/S4/200 and 37/S4/69 (Fig. 2D and 2E).

Taken together, our data shows that in the context of RNA that is highly mutated across the entire ORF, the UTRs of the MRV S gene segments are not sufficient for supporting virus rescue of those genes. However, in the same general mutational context, the reversion of 200 nts at each S gene terminus to WT sequences results in WT levels of virus rescue in a manner that does not correlate with protein expression levels of the tested segments. This supports a hypothesis where the sequences necessary and sufficient to undergo the full packaging process (assortment, assembly, replication) in the MRV S genes are located within 200 nt of the gene termini.

### 200 terminal WT nts are sufficient for packaging all S gene segments into the same replicating virus

In our previous results, 200 nts were sufficient to package each S gene segment in the context of all of the other WT gene segments. As we do not understand the extent of flexibility within the packaging process, it remained possible that the WT sequences in the other S gene segments were able to find an alternative mechanism to include the mutated segment in the recombinant viruses. To examine whether 200 nt at each termini in all four S gene segments could facilitate packaging into the same virus, we performed RG using pT7 200/S1/200, pT7 200/S2/200, pT7 200/S3/200, and pT7 200/S4/200 plasmids combined with WT M and L expressing plasmids, and measured the viral titer of the RG lysates using plaque assay. Interestingly, recombinant viruses were recovered at levels not significantly different from samples where all WT plasmids were used (Fig. 3A). Similar to results in Fig. 2, when RG was performed using all four S segments possessing wobble mutations throughout the ORF, with only WT UTRs and Stop/Start codons at each terminal end (18/S1/40, 23/S2/59, 32/S3/73, 37/S4/69) no virus was recovered (Fig. 3B). Taken together with our previous results, these findings strongly suggest that the signals required for packaging the S gene segments into a replicating virus are localized within the terminal 200 nts of each of the four S gene segments, and that favorable RNA:RNA interactions that occur during packaging between the S genes and other segments, as well as important RNA:protein interactions likely occur within and require these 200 terminal nt regions.

**Fig. 3.**
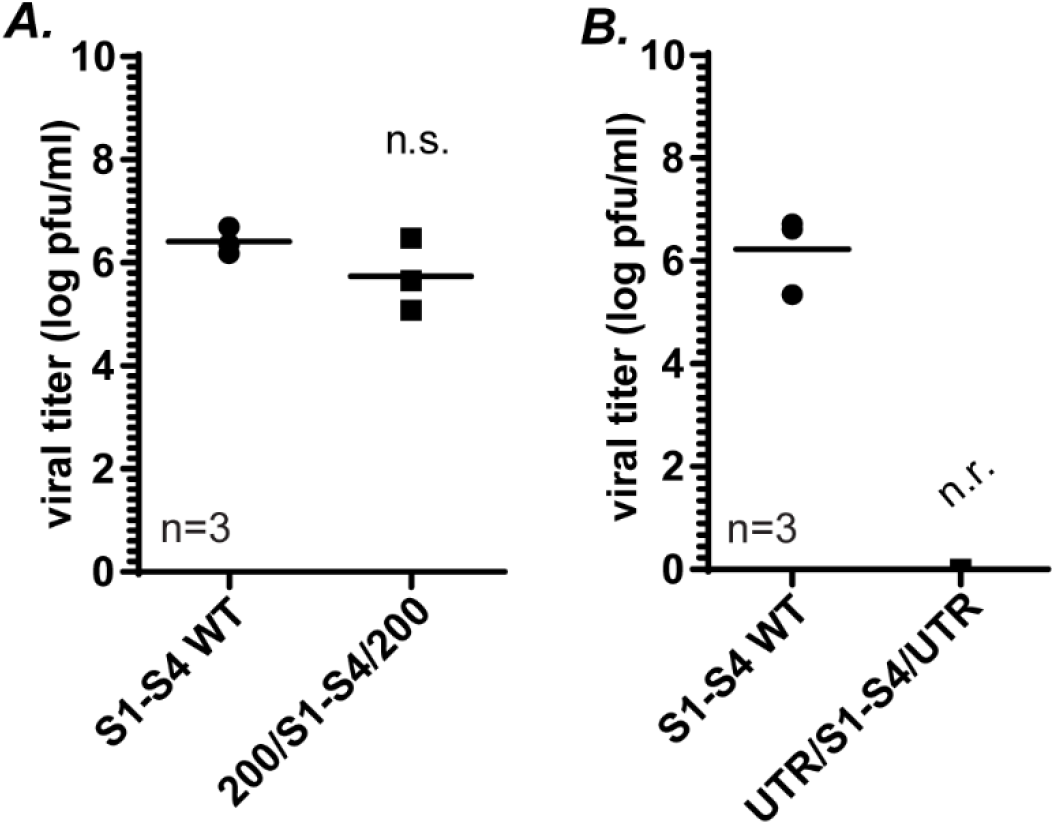
200 terminal WT nts are sufficient for packaging all S gene segments into the same replicating virus. Viral titers (log pfu/ml) of recovered recombinant viruses following RG and plaque assay using WT (S1-S4 WT) and (A) 200/S gene/200 S1,S2,S3 and S4 plasmids (200/S1- S4/200) or (B) UTR/S gene/UTR S1,S2,S3 and S4 plasmids (UTR/S1-S4/UTR) with the 6 other WT (M and L genes) plasmids. One-way ANOVA and Tukey comparisons were used for statistical analyses of data. n= independent biological replicates, n.r.=virus not recovered, n.s.=non-significant, p>0.05.

### 25 5’and 50 3’ nts are necessary for efficient S1 packaging in the W/BR assay

Our previous results identified a stretch of 200 nts at each terminus sufficient for packaging each of the S gene segments into replicating viruses. However, these nts may not be the minimal number sufficient for packaging. Our long-term goal is to better understand the MRV packaging process by identifying minimum sequences necessary and/or sufficient for packaging each individual MRV gene segment. Towards this end, we further examined the S1 gene by creating a series of plasmids where the terminal WT sequences were progressively narrowed at both the termini from 200nts to 50nts (pT7 200/S1/200, pT7 100/S1/100, pT7 50/S1/50) (Fig. 4A). These plasmids were subjected to RG assay and cell lysates were used to determine virus recovery by plaque assay. These experiments demonstrated that the 200 WT nts at the terminal ends could be narrowed to just 50 nts of WT sequence without any loss to rescue efficiency, suggesting that the 5′UTR+37 nt and the 3′UTR+13 nt are sufficient for packaging the S1 gene into a replicating virus (Fig. 4B).

**Fig. 4.**
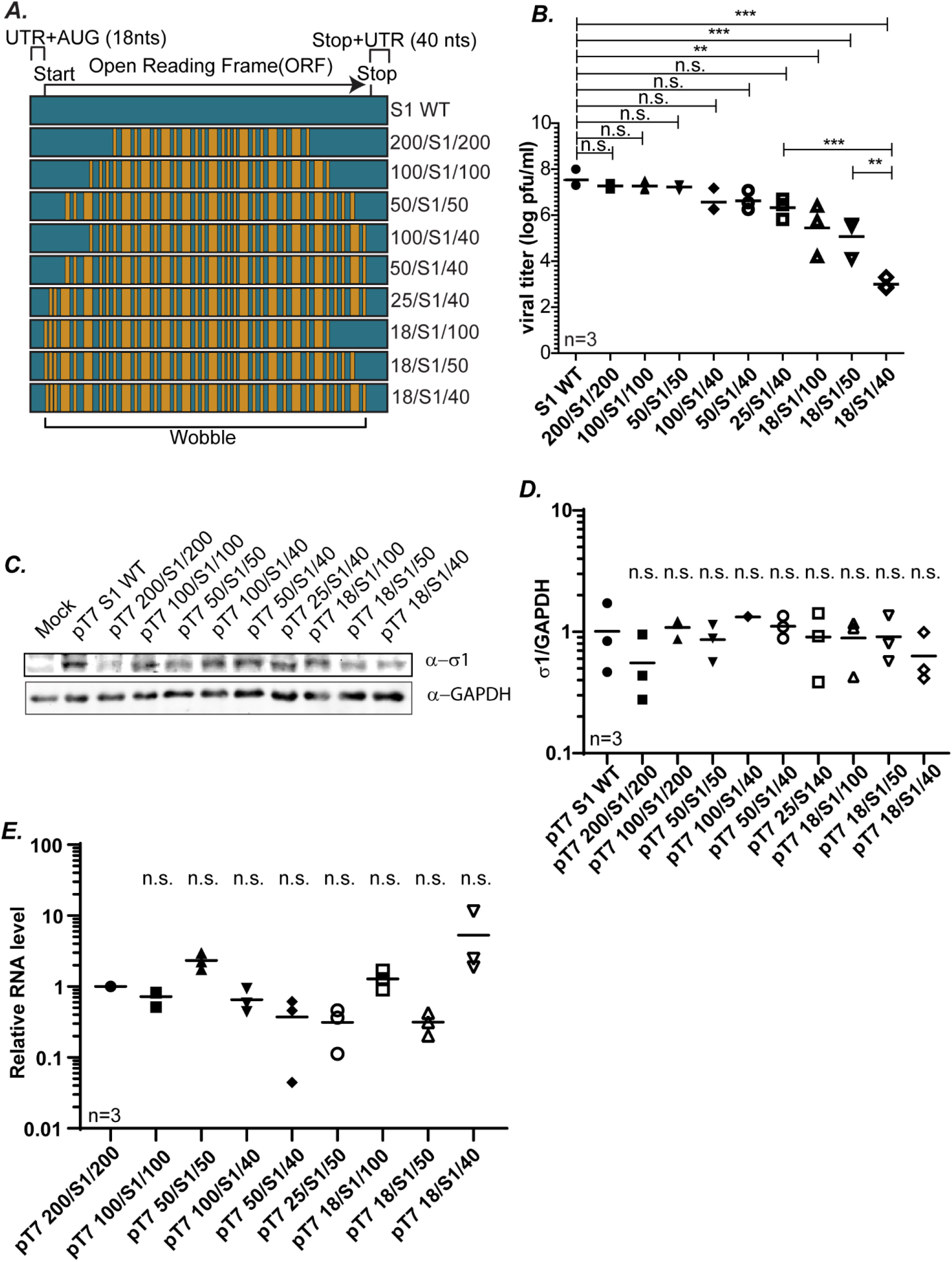
25 5’and 50 3’ nts are necessary for efficient S1 packaging in the W/BR assay. (A) Illustration of S1 gene segment mutant constructs designed to determine the minimum sequences required for packaging of the S1 gene. (B) Viral titers (log pfu/ml) determined from RG followed by plaque assay for indicated S1 mutant gene segments. (C) Representative immunoblots using σ1 antibodies showing σ1 protein expression from indicated S1 gene plasmids 24 h post- transfection. (D) Quantification of biological replicates of immunoblots in (C). σ1 protein intensities are normalized to the corresponding GAPDH loading controls. (E) Relative S1 RNA levels measured using RT-qPCR and normalized to housekeeping gene β-actin. All statistical analysis were done using one way ANOVA and Tukey multiple comparison. **p<0.01, ***p<0.001, n.s.=non-significant, p>0.05.

We were curious to determine the individual role of each terminal end in guiding the packaging process. We therefore created additional plasmids containing WT sequences of varying lengths at either the 5′ end (pT7 100/S1/40, pT7 50/S1/40, pT7 25/S1/40) or the 3′ end (pT7 18/S1/100, pT7 18/S1/50) with only WT UTR/Start/Stop sequence at the opposite end of the gene (Fig. 4A). These plasmids were subjected to RG assay and cell lysates collected and used in plaque assays to determine the impact of each mutation on virus rescue. From these experiments, we determined that with only 25 WT nts at the 5′ end (25/S1/40), there was no significant difference in the ability to recover virus compared to the fully WT S1. As demonstrated in Fig. 2, and repeated in these experiments, when a plasmid with WT sequence at the 5′ end is limited to just the UTR+Start+2 nt (18/S1/40), there is a precipitous drop of almost 4 logs in viral recovery (Fig. 4B). These findings indicate that 25 nts at the 5′ end is sufficient for packaging the S1 gene segment, and further, that sequences between nts 18 to 25 play a critical role in the packaging process. When the 3′ end contained varying stretches of WT sequences and the 5′ end was wobbled beyond the UTR/Start+2 nts (18/S1/100, 18/S1/50), there was a significant drop in viral titer of 2-3 log fold compared to WT S1 gene segment (Fig. 4B). This again reflects the relative significance of the 5′ end in the packaging process. However, the 3′ end also clearly contributes to S1 gene segment packaging, as recovery of replicating virus drops by an additional two logs in the 18/S1/40 construct, which differs by only 10 nts from 18/S1/50. Overall, our data indicate that in the context of our wobble/block replacement assay, 25 nts at the 5′end and 50 nts at the 3′ end are sufficient for packaging the S1 gene.

To determine if differences in viral rescue are due to differences in σ1 protein expression, each of the S1 mutant plasmids were transfected into BHK-T7 cells followed by immunoblotting using antibodies against σ1. These experiments revealed no significant differences in σ1expression (Fig. 4C, 4D) suggesting that differences in viral titer cannot be attributed to any difference in the translation of the S1 wobble mutant constructs. Additionally, to rule out the possibility that differences in viral recovery are due to differences in RNA availability, we measured relative RNA levels using RT-qPCR. BHK-T7 cells were transfected with the S1 mutant plasmids and 24 hours post-transfection, cells were collected and the accumulated levels of S1 RNA were measured and normalized to the housekeeping gene *β-actin* (*ACTB*). The normalized RNA levels of the S1 mutant constructs was made relative to the reference gene of 200/S1/200 for which viruses were recovered to WT S1 levels (Fig. 2A). Similar to the protein expression data, RT-qPCR showed no significant differences in relative RNA levels of the mutant S1 constructs. This suggests that differences measured in viral recovery is likely due to the effect of mutations on the process of packaging (assortment, assembly and/or replication), and not due to differences in transcription or translation of the mutant constructs.

### 5′ and 3′ UTRs are necessary for S1 gene packaging

In our previous results (Fig. 2B), we found that there is minimal rescue of recombinant virus when the S1 gene contains a wobbled ORF and WT UTRs (18/S1/40), suggesting that the UTRs are not sufficient for optimal packaging of the S1 gene. However, these experiments do not rule out the likely possibility that the UTRs play a significant role in the packaging process. To specifically address this question, we created plasmids where UTR purine nts were mutated to pyrimidines and vice versa at the 5′ UTR, 3′UTR, and both the 5′ and 3′UTR (Fig. 5A). In all the UTR mutant constructs, the 5′ GCUA and 3′ UCAUC conserved sequences that are predicted to bind the polymerase complex to guide negative-strand synthesis remained WT (27, 34). Each of these plasmids was used as a replacement for pT7 S1 WT in a RG assay, and viral titer was determined on the cell lysates by plaque assay. Regardless of whether mutations were present within the 5′UTR, 3′UTR or both the UTRs, no recombinant viruses were recovered. (Fig. 5B).

**Fig. 5.**
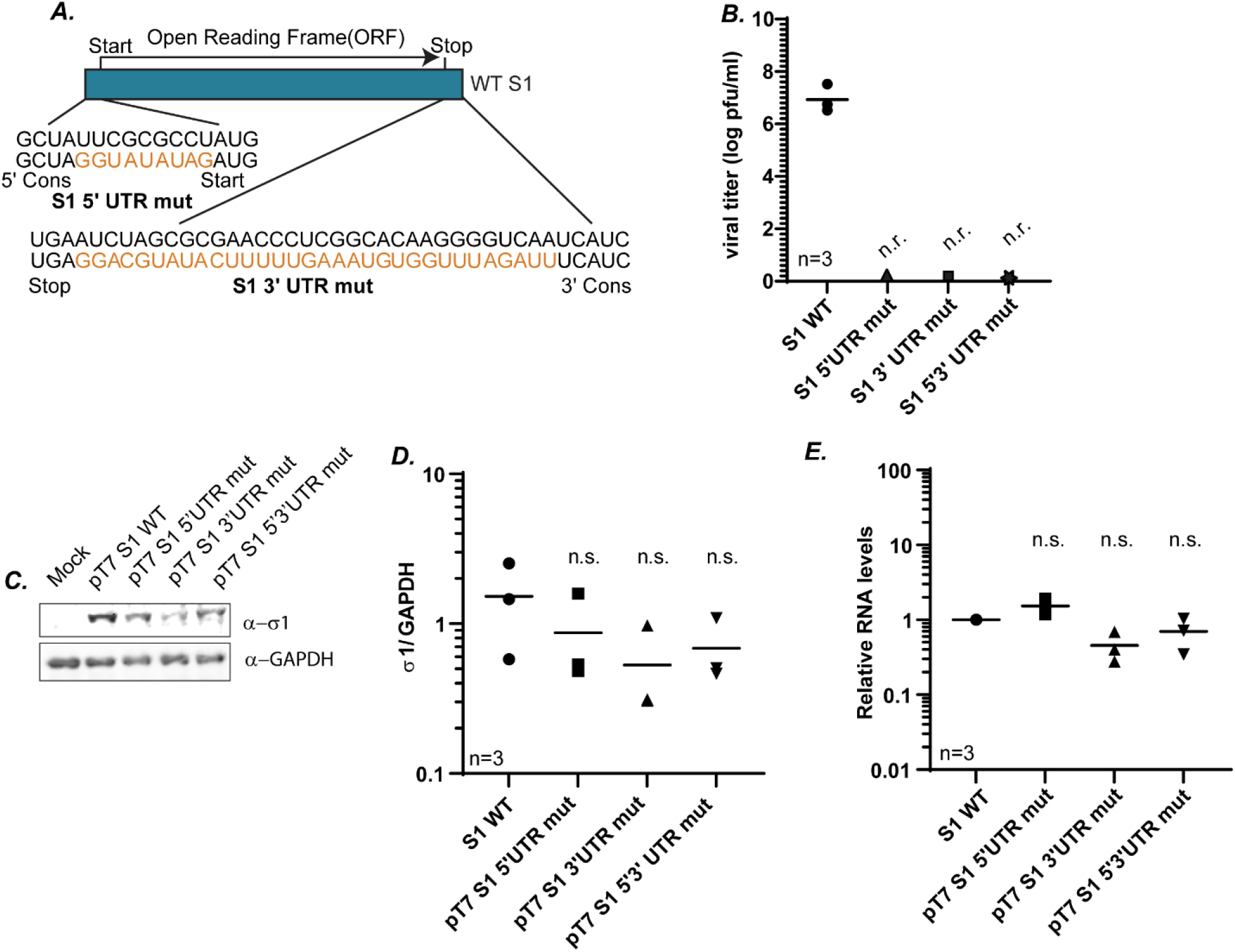
5′ and 3′ UTRs are necessary for S1 gene packaging. (A) Illustration of UTR mutations introduced into the WT S1 gene segment construct to determine the role of UTRs in S1 gene packaging. UTR mutations are shown in orange. The nt changes were introduced into the 5′UTR (S15′UTR mut), the 3′UTR (S1 3′UTR mut) or both the 5′ and 3′UTR (S1 5′3′ UTR mut). The highly conserved 5′ GCUA and 3′ UCAUC at the extreme termini were not mutated B) Viral titers (log pfu/ml) determined from RG followed by plaque assay for indicated S1 mutant gene segments. (C) Immunoblots using σ1 antibodies showing σ1 protein expression from indicated S1 gene plasmids 24 h post-transfection. (D) Quantification of biological replicates of immunoblots in (C). σ1 protein intensities are normalized to the corresponding GAPDH loading controls. (E) Relative S1 RNA levels measured using RT-qPCR and normalized to housekeeping gene β-actin. All statistical analysis were done using one way ANOVA and Tukey multiple comparison. n.r.=not recovered, Cons=conserved, n.s.=non-significant.

To rule out differences in protein expression as a factor in loss of recombinant virus rescue, pT7 5′UTR mut, pT7 3′UTR mut, and pT7 5′3′UTR mut plasmids were transfected into BHK-T7 cells, followed by immunoblotting using antibodies against σ1. There were no significant differences in protein expression between these constructs and pT7 S1 WT, suggesting the loss in virus rescue was not a result of decreased protein expression (Fig. 5C, 5D). Furthermore, there were no significant differences in gene expression of S1 from pT7 5′UTR mut, pT7 3′UTR mut, and pT7 5′3′UTR mut as compared to S1 WT as measured by RT-qPCR, suggesting that differences in S1 RNA levels between the mutant and WT plasmid constructs cannot be attributed to the lack of viral recovery. Taken together with our earlier findings, these results suggest that while the UTRs are not sufficient to support the packaging process of the S1 gene segment (Fig. 4B), they are necessary for this process as mutations within the S1 UTRs at either terminus leads to a total loss of virus recovery.

### 50 WT 5′ and 3′ terminal nts of the S1 gene are sufficient to package a non-MRV gene into infectious virions

Our previous results indicate that 25 nts at the 5′ end and 50 nts at the 3′end are sufficient for packaging the S1 gene segment into replicating viruses (Fig. 4B). However, the possibility remains that WT nt sequences remaining within the ORF after wobble mutation introduction continue to contribute to packaging in the W/BR assay. To confirm that remaining internal WT sequences do not play a role in the packaging process, we developed a novel segment incorporation assay. For this assay, plasmids were created in which the S1 ORF was replaced with the ORF for NanoLuc (NL) luciferase flanked on each end with putative S1 packaging sequences identified from our previous results. Six plasmids, which contained the NL ORF flanked on each end with S1 5′ and 3′ terminal sequences were created (pT7 S1 200/NL/200, pT7 S1 100/NL/100, pT7 S1 50/NL/50, pT7 S1 25/NL/50, pT7 S1 25/NL/40, or pT7 S1 18/NL/40) were constructed. These chimeric S1/NL plasmids were used to replace pT7 S1 WT in the RG assay. As the S1 ORF was replaced with the NL ORF, an additional plasmid (pCI-S1) which encodes the S1 ORF but does not contain the S1 5′ and 3′ UTRs and therefore cannot be packaged into recombinant viruses was included in the RG transfection to perform the function of the σ1 protein during virus recovery. As a negative control, the pCI-S1 plasmid was not included in the RG assay. RG lysates were collected at three days post-transfection because infectious viruses made in these assays will be replication-defective and will only produce one round of virus projeny. Cell lysates were collected and viruses that packaged the NL gene during the RG assay were used to infect L929 cells. Incorporation of NL was detected by measuring NL expression in the infected L929 cells (Fig. 6A). From these experiments infectious and replication-deficient virions that expressed NL luciferase were recovered when the NL ORF was flanked at each end with 200, 100, or 50 S1 nts. (Fig. 6B). This was not surprising as we also recovered viruses in our W/BR assay when these sequences were WT in an otherwise wobbled S1 ORF (Fig. 4B). Also, in line with our earlier results, we were unable to recover NL luciferase expressing virions using the pT7 18/NL/40 construct (Fig. 6B) confirming that the S1 UTRs are insufficient for packaging the gene segment. However, unlike our previous results, in these experiments we were unable to recover NL expressing viruses when pT7 25/NL/50 or pT7 25/NL/40 plasmids were used in the RG assay (Fig. 6B). This suggests that in the context of a NL ORF, 25 S1 WT nts at the 5′ end are not sufficient for packaging the segment and that WT sequences between nt 25 and nt 50 were likely contributing to rescue in the W/BR context. Each of the S1/NL/S1 chimeric plasmids were transfected individually into BHK-T7 cells and measurement of NL expression showed that they expressed similar levels of NL as compared to the positive control (pNL3.1), suggesting that differences in NL expression following viral rescue cannot be attributed to differences in protein expression (Fig. 6C).

**Fig. 6.**
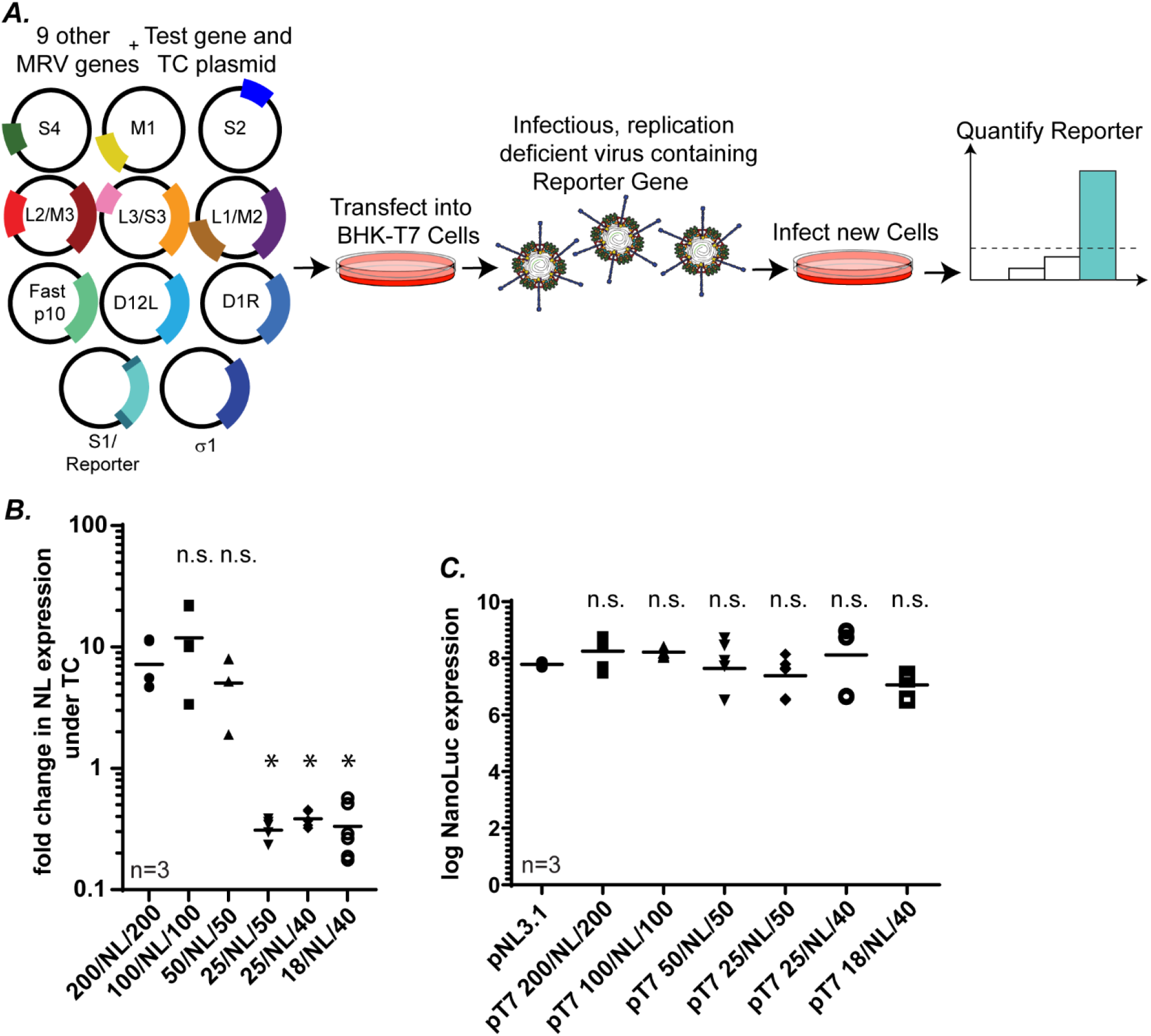
50 WT 5′ and 3′ terminal nts of the S1 gene are sufficient to package a non-MRV gene into infectious virions. (A) Schematic of the Segment Incorporation Assay. A chimeric S1 construct is designed (S1/ reporter) where the S1 ORF is replaced with a NL ORF that is flanked on both the ends with terminal S1 WT nts of varying lengths. This chimeric reporter plasmid is transfected into BHK-T7 cells with the other WT MRV gene segments and accessory plasmids, as well as a transcomplementing plasmid expressing the σ1 protein. 5 days post RG, rescued infectious virions are isolated and added onto new L929 cells. Incorporation of the chimeric gene segment is assayed by measurement of reporter gene expression (luciferase). (B) Segment incorporation assay using indicated S1/NL chimeric plasmids. (C) NL expression of control NL (pNL3.1) and indicated chimeric S1/NL plasmids measured 24 hours post transfection. All statistical analysis were done using one way ANOVA and Tukey comparison. *p<0.05, n.s.= non-significant, p>0.05).

### Sequences predicted to form an RNA panhandle structure between the 5′ and 3′ termini play a role in S1 packaging

In addition to defining S1 terminal sequences critical for packaging the S1 segment, we were also interested in examining the potential role of RNA structures formed by the interaction of the 5′ and 3′ ends in the packaging process. Our previous results provide evidence that the 5′ and 3′ terminal ends of S1 are both necessary for efficient virus rescue (compare rescue of 25/S1/40 with 18/S1/40 and 18/S1/50 with 18/S1/40 in Fig. 4B). This suggests there may be cooperation between the termini in the packaging process, which may present as direct interaction. It has been computationally predicted that MRV RNAs form a panhandle structure by the specific interaction of the 5′ and 3′ terminal nts (35). However, this panhandle structure has not been verified experimentally, and its functional significance remains unclear. To begin to explore the possibility of an interaction between the 5′ and 3′ terminal ends of S1, we folded and aligned S1 gene sequences from the three major MRV serotypes using the program Multilign (36), then manually edited the outputs to align the predicted structure (Fig. 7A,7B). From this analysis, base pairing is possible between the 5’ and 3’ ends of each serotype to form a structurally similar panhandle (Fig. 7B). Combining the *in silico* analysis with our S1 mapping data, we hypothesized that interactions between the first 25-50 and last 40-50 nts of S1 may be required for panhandle stabilization, and that perturbation of this structure may contribute to decreased mutant rescue. When analyzed using the RNA secondary structure prediction tool mfold (37), we determined that the S1 mutant constructs that were rescued from our previous results such as 25/S1/40 (Fig. 4B) are predicted to form a panhandle structure identical to the panhandle structure predicted to form for the WT S1 gene (Fig. 7B, 7C). However, the 18/S1/40 construct for which we saw a significant decrease in viral recovery (Fig. 2A, 4B) and plaque size (Fig. 2B) had a clear change in its overall panhandle architecture with shortening of the stem structure, formation of an additional stem-loop, and changes in nt base pairing (Fig. 7C).

**Fig. 7.**
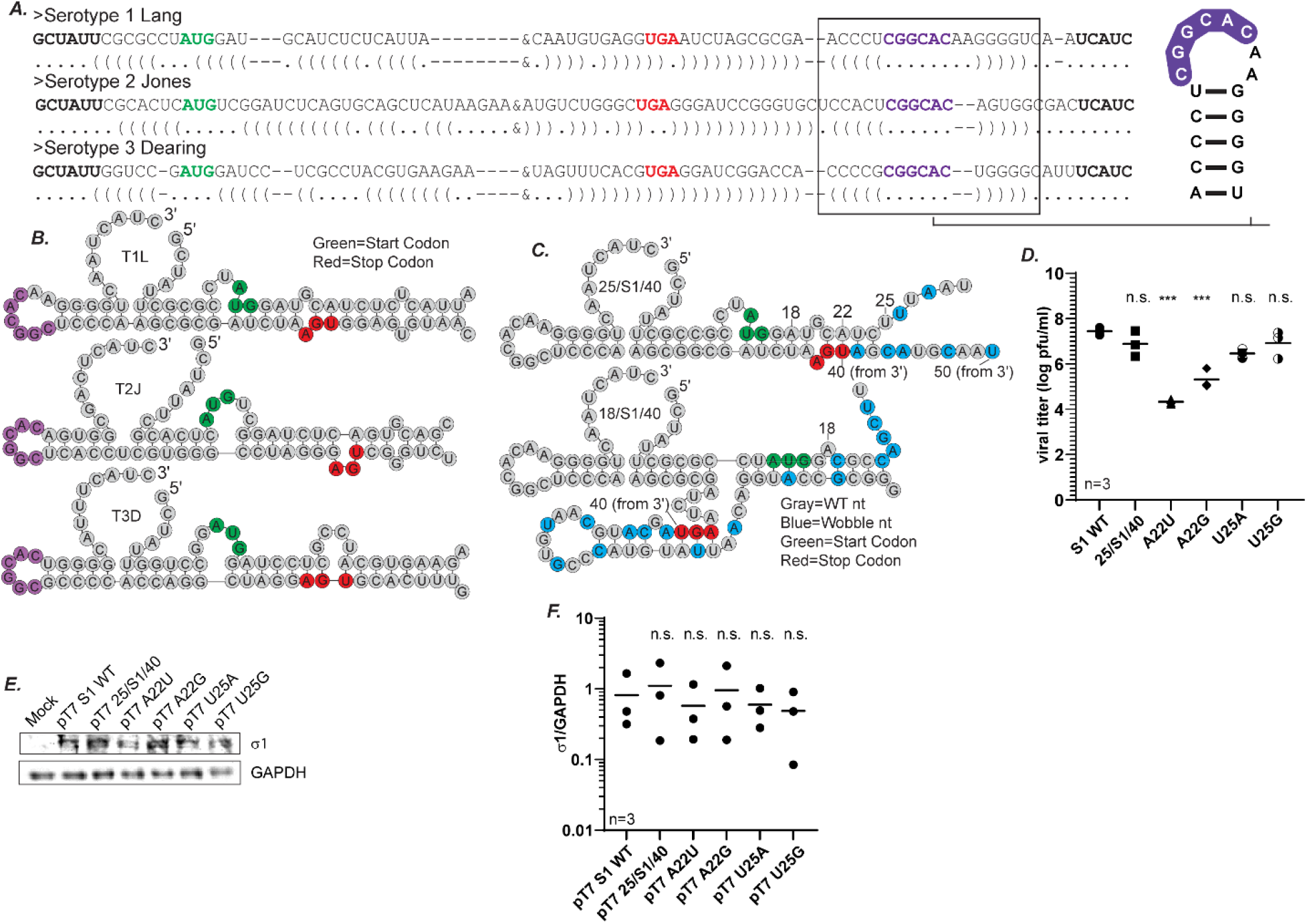
Sequences predicted to form an RNA panhandle structure between the 5′ and 3′ termini play a role in S1 packaging. (A) Dot-bracket and (B) Illustration of alignment of the terminal 5′ and 3′ end of the S1 gene from three different MRV serotypes (T1L, T2J and T3D) generated using Multialign, then manually edited to align predicted structure. A stretch of 6 conserved nts (CGGCAC) highlighted in purple form a conserved loop in the predicted panhandle structure. (C) Illustration of predicted +ssRNA secondary structures from mfold for S1 WT, 25/S1/40 and 18/S1/40. Start and stop codons are highlighted in green and red respectively. (D) Viral titers following RG as measured by plaque assay of the indicated S1 gene segment mutants. (E) Representative immunoblots using σ1 antibodies showing σ1 protein expression from indicated S1 gene plasmids 24 h post-transfection. (F) Quantification of biological replicates of immunoblots in (E). σ1 protein intensities are normalized to the corresponding GAPDH loading controls. All statistical analysis were done using one way ANOVA and Tukey comparison. ***p<0.001, n.s.=non-significant, p>0.05).

Construct pT7 25/S1/40 differs from pT7 18/S1/40 by only seven nts. To investigate the possibility that the predicted panhandle plays a role in the packaging process, we mutated specific sequences in the 25/S1/40 construct using site-directed mutagenesis. Mutation sites were specifically chosen which were predicted to either 1) play a role in forming the panhandle stem, and therefore have an impact on the formation of the predicted panhandle (nt 22), or 2) not play a role in forming the panhandle stem, and therefore have little to no impact on the predicted panhandle structure (nt 25). Two mutations each were made in nt positions 22 [pT7 25/S1/40(A22U), pT7 25/S1/40(A22G)] and position 25 [pT7 25/S1/40(U25A), pT7 25/S1/40(U25G)] taking care to maintain the WT amino acid sequence. Each mutant plasmid was then used as a replacement for pT7 S1WT in the RG assay, followed by plaque assay to measure virus recovery. As predicted, mutations at nt position 25, which does not take part in the formation of the panhandle stem in the 25/S1/40 mfold prediction, did not change viral recovery relative to 25/S1/40 (Fig. 7D). Alternatively, mutations in nt position 22 had a significant impact on viral recovery. Changing the A residue to a G led to a 1 to 1.5 log-fold reduction in viral rescue and changing the A to a U led to a 3 log-fold decrease. The difference in the efficiency of rescue between the two mutants may be due to the fact that the A to G change maintains the ability to form a G:U non canonical base pairing interaction whereas the A to U change results in a U:U in that position, which likely disrupts base pairing. Nonetheless, either mutation of this predicted base pair that is likely involved in forming the 5′/3′ panhandle of S1 leads to a drop in recovery suggesting that the sequences within this region, and potentially the panhandle structure itself, plays an important role in the packaging process. When the plasmids were transfected into BHK-T7 cells and subjected to immunoblot analysis using antibodies against σ1, there were no significant differences in protein expression (Fig. 7E, 7F), suggesting the loss in virus rescue was not a result of decreased protein expression.

### A conserved loop in the 3′ UTR plays a critical role in S1 packaging

When the full-length S1 gene sequences from the three MRV serotypes (Type 1 Lang, Type 2 Jones and Type 3 Dearing) were aligned, a stretch of 6 nts that are conserved within the 3′ UTR across all three serotypes were identified (Fig. 7A). This is unusual as the S1 gene is not well conserved compared to the other MRV gene segments (38–40). These six conserved nts form an unpaired loop in the panhandle structure which is formed by 4/5 nt pairs that contain conserved compensatory sequences across the serotypes (Fig. 8A). The complete conservation across the 3 major MRV serotypes of the predicted 6 nt loop (CGGCAC) in the 3’ UTR led us to hypothesize that this sequence/structure may also play an important role in S1 packaging. To investigate the role of this sequence, we constructed additional mutants CGGCAC to GUCACA (Loop mutant 1, LM1); CGGCAC to ACUGUG (Loop mutant 2, LM2) to change the conserved sequence without modifying the predicted panhandle/loop structure (Fig. 8A). These loop mutant plasmids were used in place of pT7 S1 WT in RG assays and cell lysates were subjected to plaque assay to determine virus titer. We were unable to rescue any replicating viruses when using these mutant plasmids, strongly suggesting the conserved sequence within the loop plays has an important function in S1 packaging (Fig. 8B). To rule out whether the introduced mutations resulted in changes in protein expression that led to a loss of viral rescue, the pT7 S1 LM1 and pT7 S1 LM2 plasmids were transfected into BHK-T7 cells, and at 24 h post transfection, immunoblotting experiments were performed. Our data showed no significant differences in σ1 protein expression (Fig. 8C, D). Additionally, when relative RNA levels were measured using RT-qPCR, there were no significant differences in gene expression of S1 between the mutant constructs and WT S1 (Fig. 8E). Altogether, these results suggest that changing six highly conserved nts that are present in an unpaired loop in an otherwise WT S1 gene leads to a complete loss of recovery of replicating viruses. These findings strongly suggest that this sequence within the unpaired loop plays a critical role in packaging the S1 gene segment.

**Fig. 8.**
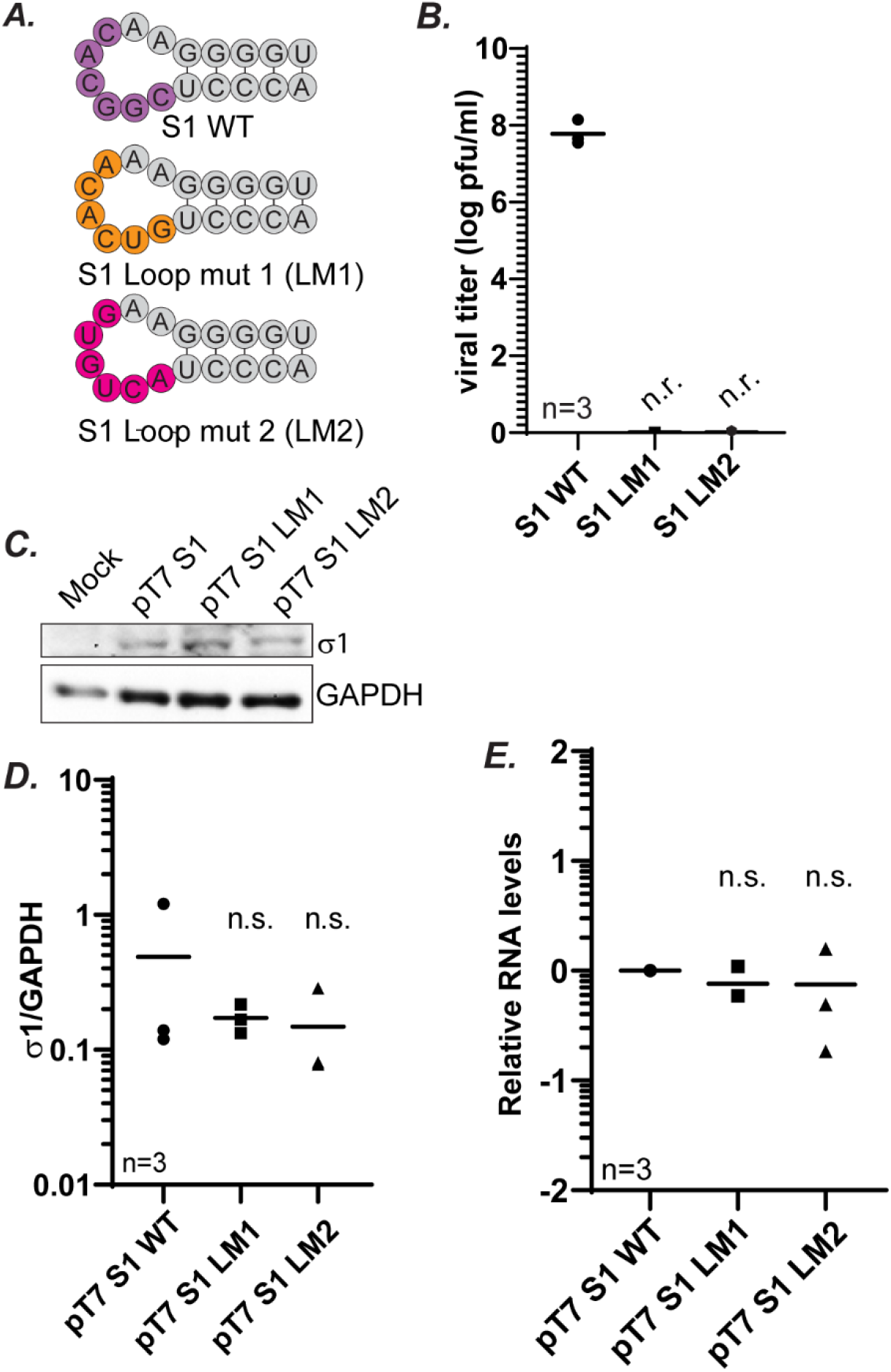
A conserved loop in the 3′ UTR plays a critical role in S1 packaging. (A) Illustration of 6 conserved nts (CGGCAC) highlighted in purple within a conserved loop in the predicted panhandle structure. Mutations that were created are indicated in orange (LM1) and pink (LM2). (B) Viral titers following RG as measured by plaque assay of the indicated S1 gene segment mutants. (C) Representative immunoblots using σ1 antibodies showing σ1 protein expression from indicated S1 gene plasmids 24 h post-transfection. (D) Quantification of biological replicates of immunoblots in (C). σ1 protein intensities are normalized to the corresponding GAPDH loading controls. (E) Relative S1 RNA levels measured using RT-qPCR and normalized to housekeeping gene β-actin. All statistical analysis were done using one way ANOVA and Tukey comparison. n.r.=not recovered, n.s.= non-significant.

### Differences in viral rescue for UTR mutants cannot be attributed to differences in expression of σ1 protein

Our previous results show that the mutations in the UTR of the S1 gene (S1 5′UTR mut, S1 3′UTR mut, S1 5′3′UTR mut, S1 LM1, S1 LM2) leads to loss of recovery of replicating virus, suggesting the UTR plays a role in the packaging process (Fig. 5B, 8B). While we provide evidence that mutation of these sequences do not result in loss of RNA or protein expression, in order to alleviate all concerns that differences in viral recovery can not be attributed to differences in expression of σ1, we performed a σ1 transcomplementation assay. In this assay we performed RG replacing pT7 S1WT with the S1 UTR mutant constructs (pT7 S1 5′UTR mut, pT7 S1 3′UTR mut, pT7 S1 5′3′UTR mut, pT7 S1 LM1, and pT7 S1 LM2). In addition a pCI-S1 plasmid which encodes only the σ1 ORF without 5′ and 3′ UTRs and will perform the function of the σ1 protein during virus recovery, was included in the RG transfection. Three days post RG under σ1 transcomplementation, viral titer was determined by plaque assay on L929 cells. Samples were additionally subjected to immunoblot using σ1-specific antibodies to measure σ1 protein levels across RG samples. Consistant with our prior experiments without σ1 transcomplementation (Fig. 5B, 8B), results from this assay show no virus rescue in the UTR mutant constructs (Fig. 9A), even in the presence of similar expression of σ1 across all the constructs tested (Fig. 9B). These results provide additional evidence that differences in viral recovery of the UTR mutants relative to WT S1 cannot be attributed to differences in expression of σ1 protein.

**Fig. 9.**
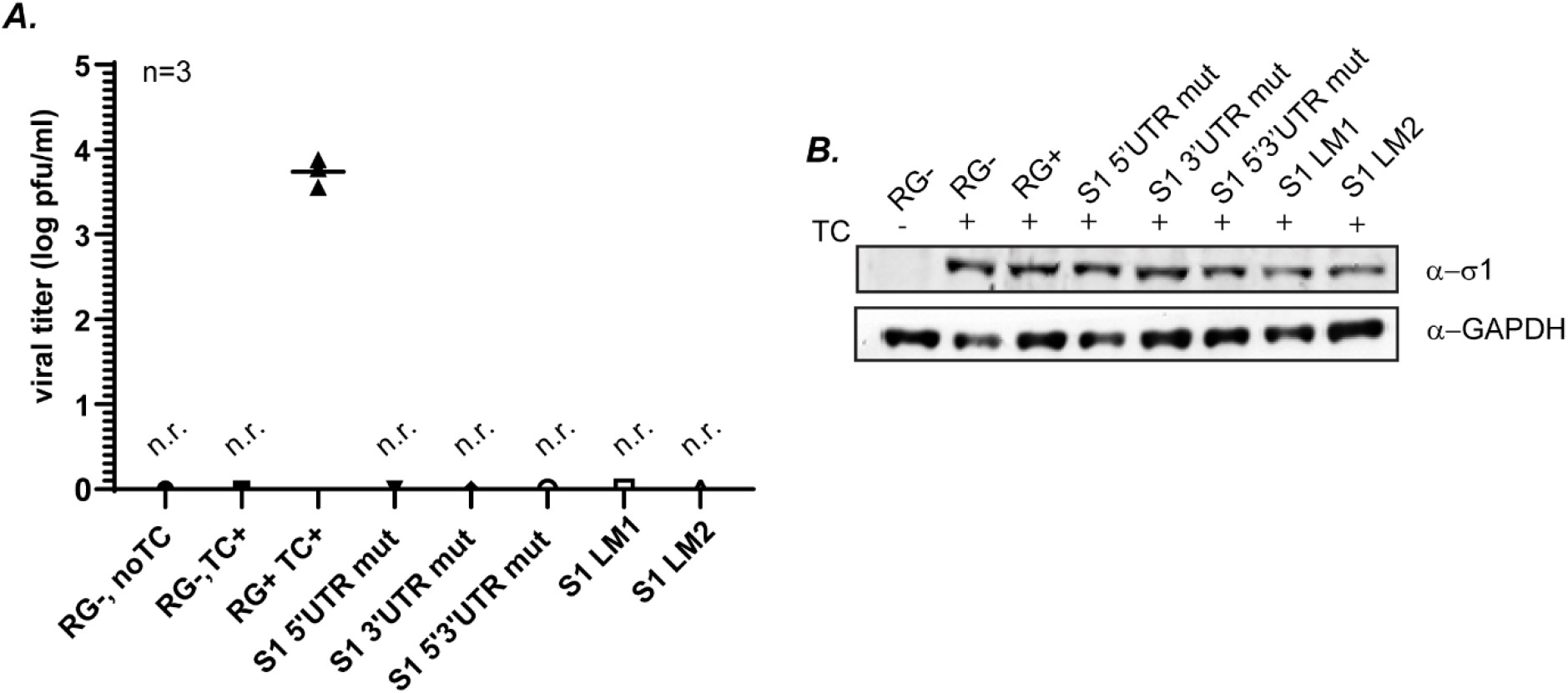
Differences in viral rescue for UTR mutants cannot be attributed to differences in translation of σ1 protein. (A) Viral titer of indicated S1 UTR mutants following RG assay transcomplemented with a plasmid expressing WT σ1 as measured by plaque assay of RG lysates on L929 cells. (B) Representative immunoblot (of 3 biological replicates) of σ1 protein measured 3 days post RG in indicated transcomplementation assays using σ1 antibodies. TC=transcomplementation, RG-, Reverse genetics negative control (no pT7-S1). RG+: Reverse genetics positive control (pT7-S1 WT), n.r.=not recovered.

## DISCUSSION

Despite decades of research, very little is understood about the mechanisms driving the packaging process of MRV. In this work, we have designed a novel assay to aid in defining viral packaging in the context of replicating viruses through introduction of wobble codons throughout the ORF. Using this approach, sequences sufficient for packaging the T1L S gene segments (S1- S4) and and the minimal nt sequences sufficient for packaging the S1 gene have been identified for the first time. In another first, we have provided experimental support that RNA secondary structure plays a critical role in the MRV packaging process, which was previously only predicted computationally (35). Furthermore, we have identified a stretch of six nts conserved across MRV serotypes that are part of an unpaired loop within the panhandle structure as critical to the packaging process. The information we have gained from our research reveals new details about the MRV packaging process and leads to many additional exciting questions to be addressed in future research.

We have identified that the packaging sequences for the S gene segments are contained within the terminal 50-200 nts of the gene inclusive of the UTRs. In future work, we would like to determine the specific functional significance of our packaging signals and clearly identify which step in the packaging process (assortment, assembly, replication, or more than one of the processes) these sequences are required. All MRV gene segments have a conserved GCUA at the 5′ UTR and a UCAUC at the 3′UTR (25) that likely functions as the binding site for the MRV RdRP protein λ3, and is essential for replication and transcription of mRNA (34). In the mutant constructs included in this study, these conserved residues were not mutated, therefore it seems less likely that λ3 binding, second-strand synthesis or mRNA synthesis have been impacted in our studies. Recent studies also suggest the polymerase of dsRNA viruses are required to bring the assortment complex to the assembling virion (22, 41). Again, because we have not altered the putative λ3 binding site in our mutants, it also seems unlikely our mutants would alter the core assembly process. For these reasons we believe the most likely explanation for the decreased/loss of virus rescue of many of these mutants is that they disrupt RNA:RNA interactions within and between gene segments and interfere with assortment of the MRV genome. Confirmation of this hypothesis will depend on the development of an MRV assortment-specific assay to demonstrate that these mutants are defective at the assortment and not some other stage of packaging.

Another question of interest is whether sequences required for packaging are located exclusively at the terminal ends of the MRV segments. Our data shows individual S gene segments, and a combination of all four S gene segments with 200 WT terminal nts flanking wobbled sequence can be packaged into replicating virus. Moreover, when just the UTRs are WT and the remaining ORF is wobbled, the S gene segments cannot be efficiently packaged. This data provides strong evidence that MRV packaging signals are localized to specific sequences/structures at the gene termini, and that there is limited flexibility within the MRV genome to compensate for mutations made within the packaging sequences. This would be in contrast to another segmented virus, Influenza A, which shows high flexibility of binding between gene segments during packaging (42). In support of this idea of limited flexibility in MRV packaging, we also show that mutations within the UTRs result in total loss of viral recovery, even though all the nts in the ORF remain WT. This suggests that disruption of these sequences, or the RNA structures they form, cannot be compensated by other sequences elsewhere within the genome. Moreover, addition of just 50 S1 terminal nts onto a non-MRV gene is sufficient to package that gene into replicating virus. Taken altogether, this data may suggest that MRV (and other dsRNA viruses) have a static RNA interaction network that is heavily sequence specific and/or structure driven. If this is the case, it seems likely that some level of flexibility must be present to account for the known property of reassortment between virus gene segments that occurs in cells infected with multiple MRV serotypes.

It is possible that MRV genome assortment is driven by a central gene segment that interacts with all other gene segments. For BTV and RV, the smallest ssRNA segment has been speculated to initiate the complex network of RNA:RNA interactions, with the UTRs of the smallest segments necessary for the process (12, 14, 43) So far with our results, we have not provided evidence that supports or refutes the possibility that S gene segments play a central role in interacting with the other gene segments. Interestingly, mutation of the conserved unpaired loop in the 3′ end of S1 resulted in complete loss of virus rescue. The loss of rescue may suggest that these sequences form a specific interaction with complementary sequences in other gene segments as part of the assortment process. In support of this hypothesis, the 3’terminal end of avian reovirus S1 ssRNA segment hybridizes with avian reovirus S4 ssRNA segment *in vitro* (44). In a scan of MRV T1L segments, we found 4 genes (S4, M2, L1, and L3) that contained exact complementary sequences which could be possible RNA:RNA binding partners to this loop within S1 during assortment (data not shown). It is also possible that all 6 nts are not involved in the RNA:RNA interaction network formed during assortment, that base pair interactions that occur between these 6 nts and another gene segment are not linear in nature, or that this sequence plays a necessary, but not sufficient role in S1 gene segment assortment or the overall packaging process. Nonetheless, if these sequences are involved in joining S1 to the assortment complex, it seems unlikely that S1 plays a central role in assortment where it recruits more than one gene segment to the complex through binding the unpaired loop. While we provide clear evidence that 200 nts at the termini of the other S gene segments are sufficient for their incorporation into the assortment complex and/or other steps in the packaging process, additional fine mapping experiments will have to be performed to clarify specific sequences/structures within the termini that are necessary for this process. Moreover, while prior studies have shown that the M and L gene segment termini are also likely to be important in packaging, future experiments in our and other laboratories will be necessary to determine the mechanism of MRV gene segment assortment.

There have been developing concerns that introducing wobble mutations into coding regions may lead to decreased levels of gene expression (45). As many of the constructs used in this study contain extensive wobble mutations within the RNA, the possibility exists that changes in RNA and protein expression may be partially responsible for changes in virus rescue. Our RT-qPCR, transcomplementation assays, and immunoblot data alleviates much of this concern as we show that levels of S RNA and σ protein expression usually are not significantly different between WT and wobble mutant constructs. Moreover levels of protein expression often do not correlate to the efficiency of viral recovery from the RG assay. For example, the pT7 200/S3/200 and pT7 32/S3/73 plasmid constructs express significantly less σNS protein than the WT pT7 S3. However, the 200/S3/200 viruses are recovered to levels not significantly different from WT S3 in RG assays. This suggests that while some of the mutations introduced in our RG constructs do result in decreased protein expression, at least in the context of the RG assay, protein expression levels are not a good measure of virus rescue efficiency. Furthermore, our results from transcomplementation assays, where we provide a plasmid expressing WT protein in RG assays, provide strong evidence that the inability to recover virus from RG assays using specific constructs is not a result of less protein expression. Moreover, other mutants used in this study (eg. Loop mutants, NL constructs) corroborated our findings in the presence of either a WT S gene or NL sequence. Therefore, we can safely conclude that introduction of wobble mutations likely impacts a step during the packaging process of the viruses and not the availability of proteins or RNA for replicating virus production.

RNA folding algorithms predict that the terminal sequences in each of the 10 MRV genes form a panhandle structure through interaction of the terminal ends (35). The mfold computational prediction for the S1 gene also suggests that the determined packaging signal sequences (5′25 and 3′50) identified in this study interact as part of the panhandle structure. Our data lend experimental proof that mutants that are predicted to modify the panhandle lead to reduced viral recovery. In addition to the panhandle, the rest of the MRV gene is also predicted to be highly structured. It is possible that the introduction of wobble mutations in the ORF alters the +ssRNA structure such that internal regions of the ORF no longer form meaningful interactions necessary for some step in the packaging process, such as interaction with the other gene segments. However, our data alleviates that concern because many of the constructs used throughout this study are wobbled across much of the gene segment, yet are rescued to WT levels, suggesting that inter- or intrasegment folding parameters needed for the packaging process are maintained. In fact, the 25/S1/40 virus, which maintains only 25 WT nt sequences at the 5′ end and 40 nts at the 3′ end, is recovered to WT virus levels even though most of the gene is wobbled. It is only when changes are made in the predicted panhandle structure that viral recovery is negatively impacted, further showing the importance of the panhandle structure or sequences contained within the panhandle region. Although our data suggest that at least for the S segments RNA sequences or structure internal to the terminal 200 nts is not necessary for packaging of these segments into the MRV virion, we have not yet investigated the larger M or L segments and it remains quite possible that RNA structure beyond the panhandles, both within and between gene segments plays an important role in MRV packaging.

## MATERIALS AND METHODS

### Cells

BHK-T7 cells expressing the T7 polymerase (46) were maintained in Dulbecco′s modified Eagle′s medium (DMEM) (Invitrogen life technologies) supplemented with 10% fetal bovine serum (Atlanta Biologicals), Penicillin (100IU/ml)–Streptomycin (100 μg/ml) solution (Mediatech), and 1% MEM nonessential amino acids solution (HyClone). To maintain the expression of T7 polymerase, 1 mg/ml of G418 (Alexis Biochemical) was added after every four passages. Spinner adapted L929 cells were maintained in Joklik modified minimum essential medium (Sigma-Aldrich) supplemented with 2% fetal bovine serum, 2% bovine calf serum (HyClone), 2 mM L-glutamine (Mediatech), and penicillin (100 IU/ml)–streptomycin (100 μg/ml) solution.

### Plasmid construction

The ten (47) (ID: 33295, 33294, 33293, 33292, 33291, 33290, 33289, 33288, 33287, 33286) and four (48) (ID: 33306, 33305, 33304, 33303) reverse genetics (RG) system plasmids for T1L, along with plasmids encoding the vaccinia virus capping enzymes pCAG D1R (ID:89160) and pCAG D12L(ID:89161) and a Nelson Bay Virus syncytia inducing plasmid pCAG Fast p10 (ID:89152) were purchased from Addgene. pCI-S2 (T1L) was previously described (49). pCI-S1 is a plasmid expressing Type 3 Dearing σ1, and was a kind gift from Dr. Max Nibert. A pCI-Rz plasmid was created by PCR amplification of the Hepatitis δ ribozyme sequence and ligation into the multiple cloning site of the pCI-Neo plasmid using 5′ SacI/3′SmaI restriction endonuclease sites. WT T1L S gene segment plasmids (pT7 S1WT, pT7 S2WT, pT7 S3WT, pT7 S4WT) were generated by RT-PCR amplification of viral S1, S2, S3, and S4 gene segments respectively from purified T1L virus with primers that contain a 5′ SacI cleavage site upstream of a T7 promoter sequence. The 5′ SacI-T7-S gene segment 3′ PCR product was then ligated into the pCI-Rz plasmid using the following cleavage sites: S1: 5′SacI/3′XhoI (partial restriction digestion as S1 ORF has an internal SacI site), S2: 5′SacI/3′AvaI, S3: 5′SacI/3′NheI, S4: 5′SacI/3′XbaI.

The mutated S gene segments were synthesized as double-stranded (ds) DNA sequences (gBlocks™) by Integrated DNA Technologies with restriction endonuclease cleavage sites at the 5′ and 3′ end of the dsDNA. The gBlocks were digested and cloned into the parent gene segment plasmids (pT7 S1WT, pT7 S2WT, pT7 S3WT, pT7 S4WT) following the NEB Cloner workflow. Specific mutant constructs were subcloned from the mutants using appropriate restriction digestion sites or using the site-directed mutagenesis kit (Quick Change II Site directed mutagenesis kit, Agilent Technologies) following manufacturers protocol. The 5′ and 3′ restriction endonuclease sites, the vector backbones used, primer sequences used in site directed mutagenesis for making the mutant S gene segment constructs are listed in Table 1.

**Table 1.**
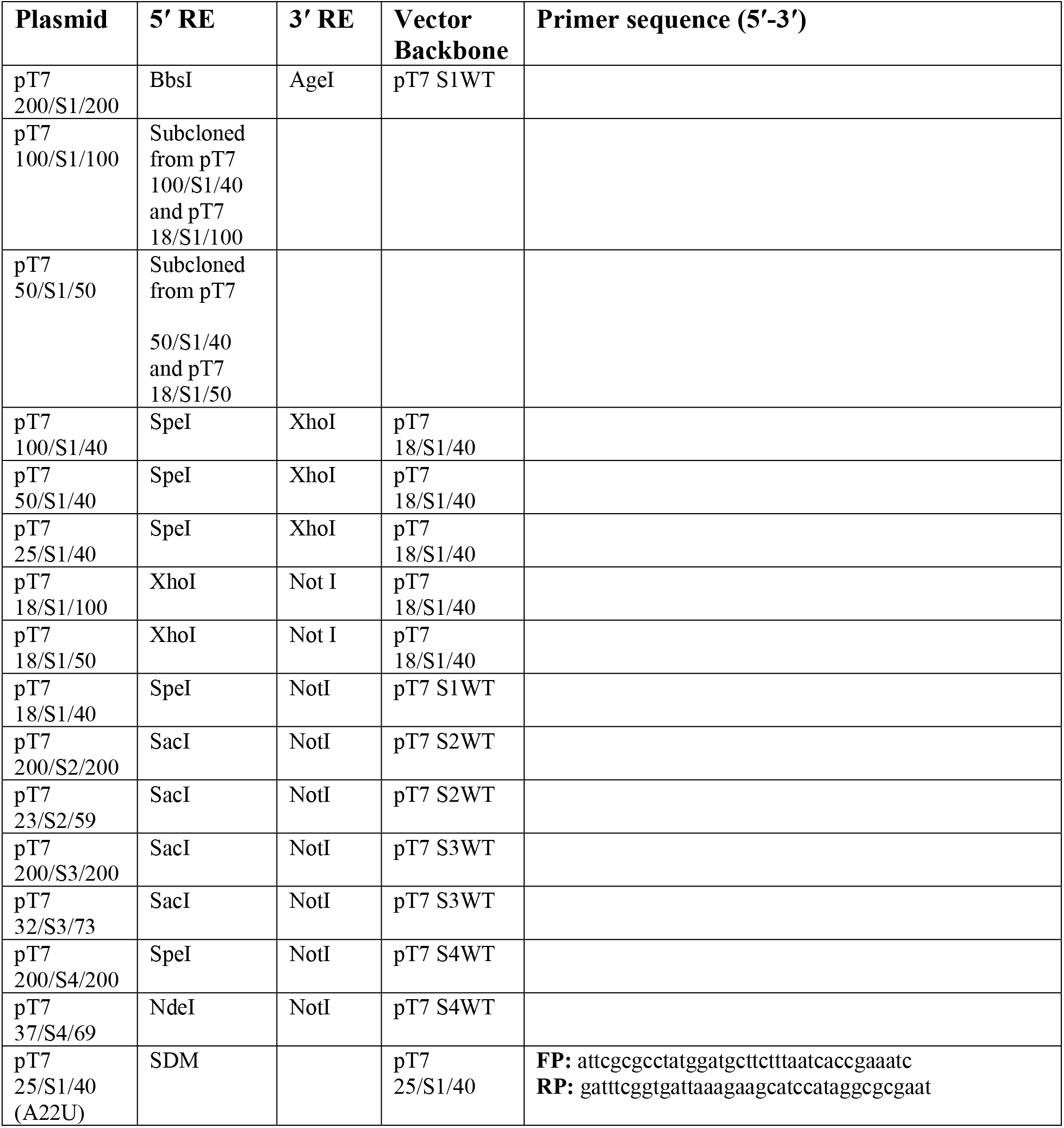
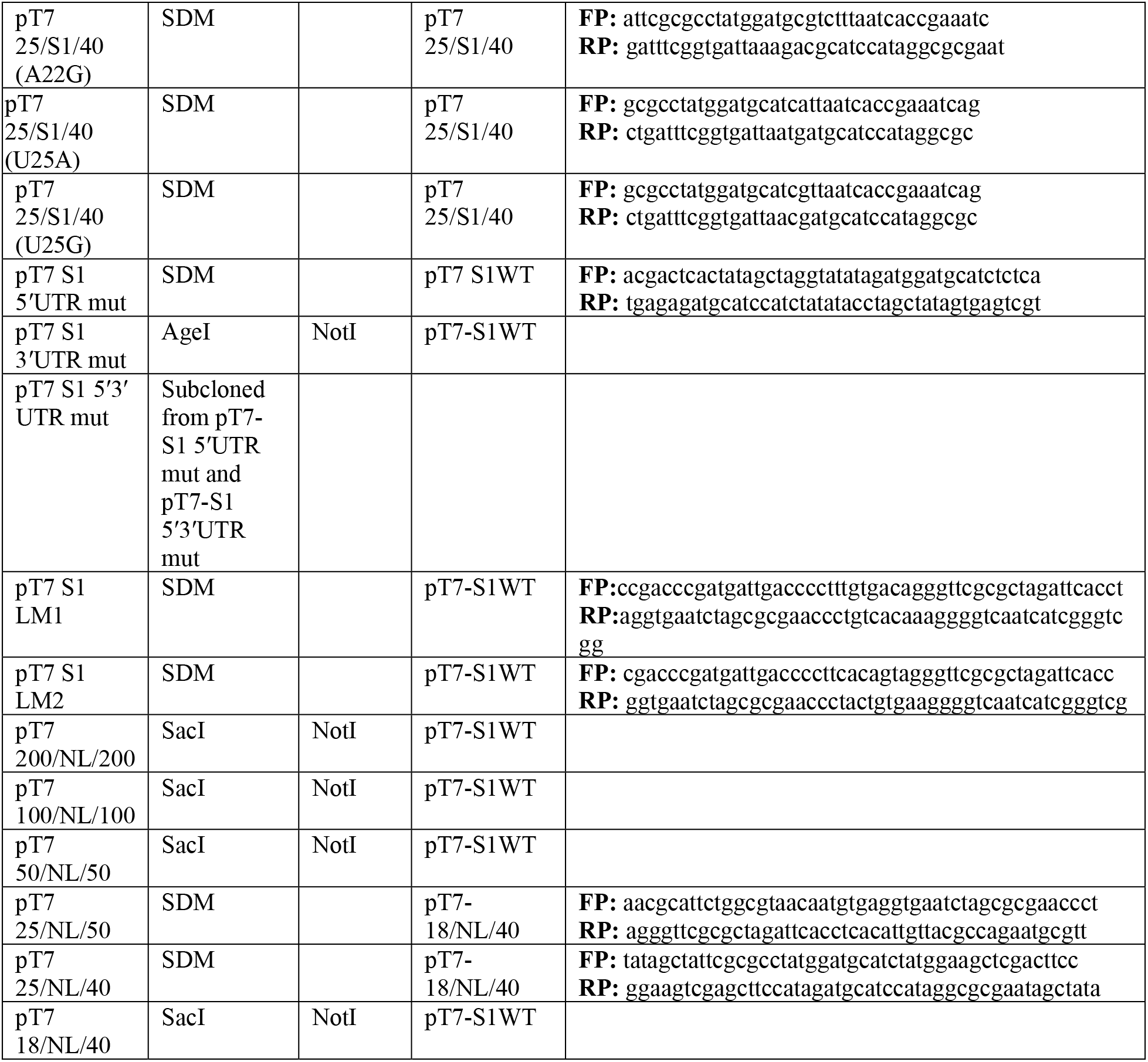
Mutant S gene segment plasmid design. Table of the mutant S gene segment plasmid constructs, with the restriction endonuclease cleavage sites used at the 5′ end (5′ RE) and the 3′ end (3′ RE) along with the vector backbone used for cloning in the insert, and the primers (Forward primer: FP and Reverse primer: RP) used during site directed mutagenesis (SDM).

### Infections and transfections

The laboratory stock of T1L is derived from the lab of B.N Fields. Viruses were purified using a method described previously (50) using Vertrel-XF (Miller Stephenson Chemical company, Catalog: NC9715008) instead of freon (51). Multiplicity of infections calculations for T1L coinfections in the Segment Incorporation Assay were based on cell infectivity unit (CIU) measurement as described previously (52–54). Transfections were carried out in Opti-MEM™ (Thermo Fisher Scientific) using TransIT-LT1 transfection reagent (Mirus, Cat: MIR 2306) following the manufacturer’s protocol with a TransTt-LT1(μl): plasmid(μg) ratio of 3:1.

### Antibodies

σ2 rabbit polyclonal antiserum against heat inactivated core (55), rabbit polyclonal σNS antiserum (56), and mouse monoclonal antibodies against σ3 (4F2, 5C3, 10C1) (57) were used for immunoblot assays. Rabbit (σ1) VU304 antisera raised against the T1L σ1 head domain was a generous gift from the Dermody lab, University of Pittsburgh. Mouse monoclonal GAPDH (Thermo Scientific^TM^, Cat: MA5-15738) and rabbit α-TUBULIN (Cell Signaling Technologies™, Cat: 50-190-297) were used as loading controls in immunoblot assays.

### Wobble/Block Replacement (W/BR) assay

BHK-T7 cells were plated overnight at 2x10^5^ in a 12 well dish. 0.4 μg of plasmids from the MRV four and ten plasmid RG system (47, 48) were transfected to express the nine WT MRV gene segments. 0.4 μg of the mutant gene segment was transfected along with the above plasmids to provide the tenth segment.

A modification was made from the original four plasmid RG system protocol (47) by the addition of plasmids expressing vaccinia virus capping enzymes pCAG D1R (0.2 μg), pCAGD12L (0.2 μg) and Nelson Bay Virus fusion associated transmembrane protein pCAGFast p10 (0.005 μg) that have been previously reported to enhance viral recovery in RG by almost 1,150 fold (58). All RG transfections were allowed to progress for 5 days unless otherwise mentioned, after which cells were subjected to three freeze-thaw cycles to recover virus.

### Plaque assay

Viral titers of replicating viruses were determined using standard plaque assay in L929 cells (50). L929 cells were plated in 6 well plates at 1.5x10^6^ per well. After overnight incubation, viral samples were serially diluted in Phosphate Buffered Saline (PBS) MgCl_2_ [0.2 mM MgCl_2_ supplemented to 1X PBS (137 mM NaCl, 3 mM KCl, 8 mM Na_2_HPO_4_, 1.5 mM KH_2_PO_4_, pH 7.4)]. Viral dilutions were added onto the L929 monolayers at room temperature for viral attachment for 1 hour. Post-viral attachment, an overlay media consisting of 2X Serum Free Medium 199 (GibcoTM) supplemented with 1X Penicillin Streptomycin and 1X L-glutamate was added along with 2 mg/ml Trypsin (final concentration: 80μg/ml), 2% Agar (1% final concentration) and 5% Sodium Bicarbonate (0.15% final concentration). Viral titers were determined by counting plaques three days post overlay addition.

### Crystal Violet staining of plaques

After determination of viral titer, 8% paraformaldehyde was added to each well to fix the L929 cells, and left to incubate overnight. The next day, overlay media and paraformaldehyde were gently removed and cells were washed once with 1XPBS. 1% Crystal Violet in 20% ethanol was added and incubated at room temperature for 15-30 min to stain the fixed L929 cells, after which the staining solution was removed and left at room temperature for 1 min. Plaque images were taken using an iPhone 13 ProMax camera set-up that maintained the plates at the same distance from the camera.

### Viral RNA extraction and sequencing

To extract viral RNA of recovered viruses from RG, individual plaques, picked using disposable glass Pasteur pipettes (Fisherbrand), were added to L929 or Vero cells in a T75 flask and passaged twice. Viral RNA was extracted using TRIzol LS (Life Technologies) following manufacturer′s instructions. Approximately 1μg of total RNA was reverse transcribed using SuperScript IV (Invitrogen Life Technologies) as per manufacturer′s protocol. cDNA was PCR amplified using gene specific primers to produce full-length gene segments, and the entire genes were sequenced using gene specific primers.

### RT-qPCR assay

5x10^5^ BHK-T7 cells were plated in 6 well dishes and after overnight incubation, transfected with 3μg of total plasmid using TransIT-LT1 transfection reagent (Mirus, Cat: MIR 2306) following the manufacturer’s protocol with a TransIT-LT1 (μl): plasmid (μg) ratio of 3:1. 24 hours post-transfection, cells were collected in PBS and total RNA was extracted using PureLink RNA miniprep kit (Invitrogen Life Technologies) following manufacturer’s protocols. Total RNA was then digested using DNase I for 2h at 37°C (New England Biolabs) to remove residual plasmid DNA and then purified using Monarch RNA cleanup kit (New England Biolabs). RNA was reverse transcribed using SuperScript IV (Invitrogen Life Technologies) or qScript Ultra Flex Kits (Quantabio), using gene specific primers (Integrated DNA Technologies). The real time PCR was performed using PowerTrack SYBR Green Master Mix (Applied Biosystems) and SYBR Green detection was done using QuantStudio3^TM^ Real-Time PCR system. The Real-Time PCR was done using standard cycling conditions [Enzyme activation: 95°C for 2 min, (Denaturation: 95°C for 15s, Anneal/Extend: 60°C for 60s) for 40 cycles]. Melt curve analysis was done at 95°C for 15s, followed by 60°C for 60s. The dissociation analysis was done at 95°C for 15s. Data analysis was done using Design and Analysis software 2.6.0 (Applied Biosystems). The data is reported using the 2^-ΔΔ*CT*^ method of relative quantification (59). Briefly the C_T_ value of the S segment PCR was subtracted from the C_T_ value of the housekeeping gene PCR to generate a normalized C_T_ value for the S segment. This normalized C_T_ value was then subtracted from the normalized C_T_ value of the reference gene to generate a ΔΔ*C_T_* value. This ΔΔ*C_T_* value was then used in the formula 2^-ΔΔ*CT*^ and compared to the reference gene for reporting the relative RNA level.

### Segment Incorporation Assay

Chimeric S1 NL plasmids and the pCI-Neo S1 plasmid were used to replace pT7-S1 in the RG assay. Two days post RG assay, media was removed, samples were washed 3 times with 1X PBS and then refed with fresh DMEM to remove untransfected plasmids from the samples. The RG reaction was allowed to proceed an additional 3 days, after which samples were subjected to three freeze-thaw cycles to release virions from cells. The cells debris was removed after centrifugation at 15000g for 5 mins and cell supernatant containing the released virions were subjected to 5μl DNase I (Zymo research) for 1 hour to remove remaining untransfected chimeric NL plasmid. The DNase I treated samples were then co-infected with WT T1L into L929 cells plated in 12 well dishes at 2x10^5^ cells per well. Twenty-four hours post-infection, cells were gently washed and collected in sterile 1XPBS. The cells were treated with NanoGlo Luciferase reagent (Promega) according to manufacturer′s protocol and Luciferase activity was measured using a Glomax Multi Detection Plate reader on a 96 well plate.

### SDS-PAGE and Immunoblot

5x10^5^ BHK-T7 cells were plated overnight on 60mm dishes and transfected with 10μg of plasmids. Twenty-four hours post-transfection, cells were collected in ice cold 1XPBS and lysed with RAF buffer (20 mM Tris [pH 8.0], 137 mM NaCl, 10% glycerol, 1% NP-40) (60) for 1 hour on ice. Whole cell lysates were resuspended in 2X Laemeli loading dye (125 mM Tris-HCl [pH 6.8], 200 mM dithiothreitol [DTT], 4% sodium dodecyl sulfate [SDS], 0.2% bromophenol blue, 20% glycerol), and subjected to sodium dodecyl sulfate-polyacrylamide gel electrophoresis (SDS-PAGE) for protein separation. Membranes were blocked using 5% Milk in TBS-T (20 mM Tris, 137 mM NaCl [pH 7.6]) with 0.1% Tween 20 and treated with primary and secondary antibodies for 18 h and 2 h respectively with 3x15 minute TBS-T washes after each antibody incubation. Membranes were treated with NovaLume Atto Chemiluminescent Substrate AP (Novus Biologicals) using manufacturers protocol and imaged on a ChemiDoc XRS Imaging System with Quantity One imaging software (Bio-Rad Laboratories).

### Protein Transcomplementation Assay

0.4 μg S segment mutant constructs (pT7 23/S2/59, pT7 S1 5′UTR mutant, pT7 S1 3′UTR mutant, pT7 S1 5′3′ UTR mutant, pT7 S1 LM1, pT7 S1 LM2) were subjected to the RG procedure as described above. σ protein was provided by transfecting 0.4 μg of pCI-S1 (for S1 5′UTR mutant, S1 3′UTR mutant, S1 5′3′ UTR mutant, S1 LM1, S1 LM2 RG) or 0.4 μg of pCI-S2 (for 23/S2/59 RG). Three days post RG, cells were subjected to three freeze thaw cycles. Virus replication was measured by plaque assay on L929 cells and σ protein expression was confirmed by subjecting the RG samples to SDS-PAGE and immunoblotting.

### Figures, graphs and statistical analysis

All graphs and statistical measurements were performed in Graph Pad Prism Version 9 (Graphpad Software). Figures were generated using Adobe Photoshop 2021 and Adobe Illustrator 2021 (Adobe).

## ACKNOWLEDGMENTS

This publication is supported by funding from National Institute of Health (NIH) R15CA202984 and Iowa State University College of Veterinary Medicine Seed Grants.

